# Expanding the repertoire of the plant infecting ophioviruses

**DOI:** 10.1101/2023.01.27.525910

**Authors:** Humberto Debat, María Laura García, Nicolás Bejerman

## Abstract

Ophioviruses (genus *Ophiovirus*, family *Aspiviridae*) are plant-infecting viruses with non-enveloped, filamentous, naked nucleocapsid virions. Members of genus *Ophiovirus* have a segmented single-stranded negative-sense RNA genome (ca. 11.3-12.5 kb), encompassing three or four linear segments. These segments encode in total four to seven proteins in sense and antisense orientation, both in the viral and complementary strands. The genus *Ophiovirus* includes seven species with viruses infecting both monocots and dicots, mostly trees, shrubs and some ornamentals. From a genomic perspective, as of today, there are complete genomes available for only four species. Here, by exploring large metatranscriptomics publicly available datasets, we report the identification and molecular characterization of 33 novel viruses with genetic and evolutionary cues of ophioviruses. Genetic distance and evolutionary insights suggest that all the detected viruses could correspond to members of novel species, which expand ca. 4.5-fold the current diversity of ophioviruses. The detected viruses increase the tentative host range of ophioviruses for the first time to mosses, liverwort and ferns. In addition, viruses were linked to several *Asteraceae, Orchidaceae* and *Poaceae* crops/ornamental plants. Phylogenetic analyses showed a novel clade of mosses, liverworts and fern ophioviruses, characterized by long branches suggesting still plenty unsampled hidden diversity within the genus. This study represents a significant expansion of genomics of ophioviruses, opening the grounds to future works on the molecular and evolutionary peculiarity of this virus genus.

## 1. Introduction

A vast number of viruses are being discovered in this new metagenomic era, revealing a multifaceted and diverse evolutionary landscape of replicating entities and the complexities associated to their arduous classification (Koonin et al., 2021). Several strategies to lever this dynamic growing wide-ranging assemblage of viruses has led to an initial comprehensive proposal to generate a virus world megataxonomy (Koonin et al., 2020). Even though extensive broad efforts to characterize the virus share of the biosphere, only an infinitesimal portion, which probably embodies less than one percent of the virosphere appears to be characterized so far (Geoghegan and Holmes, 2017). Consequently, our knowledge about the massive global virome, with its outstanding diversity and including every prospective host organism assessed so far is scarce (Dolja et al., 2020; Edgar et al., 2022; Mifsud et al., 2022). Data mining of publically available transcriptome datasets derived from High-Throughput Sequencing (HTS) has become an efficient and inexpensive strategy to render the hidden diversity of the plant virosphere (Edgar et al., 2022; Bejerman et al., 2020). Data-driven virus discovery emerges in the context of a massive number of open datasets on the Sequence Read Archive (SRA) of the National Center for Biotechnology Information (NCBI). This wonderful reserve of sequences, which is growing at an exceptional rate, represents a substantial (but still limited and biased) portion of all the organisms that populate our world, which converts the NCBI-SRA database an efficient and cost-effective resource to identify novel viruses (Lauber and Seitz, 2022). From a virus taxonomy perspective, a consensus statement has defined that viruses that are known only from metagenomic data can, and should be, incorporated into the official classification scheme of the International Committee on Taxonomy of Viruses (ICTV) (Simmonds et al., 2017).

Ophioviruses (genus *Ophiovirus*, family *Aspiviridae*) are plant-infecting viruses with non-enveloped, filamentous, naked nucleocapsid virions. Members of genus *Ophiovirus* have a segmented single-stranded negative, and possible ambisense RNA genome, encompassing three or four linear segments (in total ca. 11.3-12.5 kb) (Garcia et al., 2017). These segments encode four to seven proteins in sense and antisense orientation, both in the viral and complementary strands (Garcia et al., 2017). The genus *Ophiovirus* includes seven recognized species with viruses infecting both monocots and dicots, mostly trees, shrubs and some ornamentals, and four out of these seven species are reported to be transmitted via soil-borne fungus of the genus *Olpidium* spp (Garcia et al., 2017). From a genomic perspective, as of today, there are complete genomes available for only four of those seven members species. In the context of a systematic expansion of virus discovery favored by the extensive use of HTS, a plethora of novel viruses of many families from diverse plants have been described. Nevertheless, to our knowledge, the diversity of ophioviruses appears to be stagnated, with no new ophiovirus species recognized by the ICTV since 2015. Two recent works have described the complete genome of a novel proposed ophiovirus associated to carrot, carrot ophiovirus 1 (CaOV1) (Fox et al., 2022) and other found in pepper, pepper chlorosis-associated virus (PCaV) (Shimomoto et al., 2023). In addition, the segment that encodes the capsid protein (CP) of a putative novel ophiovirus was assembled from transcriptomic data of *Dactylorhiza hatagirea* (Sidharthan et al., 2021).

This is the first study oriented to identify and characterize ophioviruses sequences that are hidden in publicly available metatranscriptomic data, which resulted in the identification and characterization of 33 novel tentative ophioviruses. Our findings significantly expand the *statu quo* of genomics of ophioviruses, opening the grounds to future works on the molecular and evolutionary peculiarities of this virus genus and the *Aspiviridae* family.

## 2. Material and Methods

### 2.1 Identification of Ophiovirus sequences from public plant RNA-seq datasets

Two strategies were used to detect ophiovirus sequences: 1) Assembled and Raw sequence data corresponding to the 1K study (Leebens-Mack et al., 2019) was explored by tBlastn searches (E-value < 1e^-5^) for ophiovirus sequences using the NCBI-refseq proteins of ophioviruses at the 1KP:BLAST tool (https://db.cngb.org/onekp) and hits were curated with the raw SRA data retrieved from the NCBI BioProject PRJEB4922. 2) The Serratus database was analyzed, employing the serratus explorer tool (Edgar et al., 2022) using as query the predicted RNA dependent RNA polymerase protein (RdRP) of ophioviruses available at the NCBI-refseq database. The SRA libraries that matched the query sequences (alignment identity > 45%; score > 10) were further explored in detail.

### 2.2 Sequence assembly and virus identification

Virus discovery was implemented as described elsewhere (Bejerman et al., 2022; Bejerman & Debat., 2022). In brief, the raw nucleotide sequence reads from each SRA experiment that matched que query sequences in both the 1k and Serratus platforms were downloaded from their associated NCBI BioProjects (**Table 1**). The datasets were pre-processed by trimming and filtering with the Trimmomatic tool as implemented in http://www.usadellab.org/cms/?page=trimmomatic. The resulting reads were assembled *de novo* with rnaSPAdes using standard parameters on the Galaxy server (https://usegalaxy.org/). The transcripts obtained from *de novo* transcriptome assembly were subjected to bulk local BLASTX searches (E-value < 1e^-5^) against ophiovirus refseq protein sequences available at https://www.ncbi.nlm.nih.gov/protein?term=txid88129[Organism]. The resulting viral sequence hits of each dataset were explored in detail. Tentative virus-like contigs were curated (extended and/or confirmed) by iterative mapping of each SRA library’s filtered reads. This strategy is used to extract a subset of reads related to the query contig, use the retrieved reads from each mapping to extend the contig and then repeat the process iteratively using as query the extended sequence. The extended and polished transcripts were reassembled using Geneious v8.1.9 (Biomatters Ltd.) alignment tool with high sensitivity parameters.

### 2.3 Bioinformatics tools and analyses

#### 2.3.1 Sequence analyses

ORFs were predicted with ORFfinder (minimal ORF length 150 nt, genetic code 1, https://www.ncbi.nlm.nih.gov/orffinder/), functional domains and architecture of translated gene products were determined using InterPro (https://www.ebi.ac.uk/interpro/search/sequence-search) and the NCBI Conserved domain database - CDD v3.20 (https://www.ncbi.nlm.nih.gov/Structure/cdd/wrpsb.cgi) with evalue = 0.01. Further, HHPred and HHBlits as implemented in https://toolkit.tuebingen.mpg.de/#/tools/ were used to complement annotation of divergent predicted proteins by hidden Markov models. Transmembrane domains were predicted using the TMHMM version 2.0 tool (http://www.cbs.dtu.dk/services/TMHMM/). The predicted proteins were then subjected to NCBI-BLASTP searches against the non-redundant protein sequences (nr) database to filter out any virus-like sequences that did not show as best hit an ophiovirus protein.

#### 2.3.2 Pairwise sequence identity

Percentage amino acid (aa) sequence identities of the predicted CP protein of the ophioviruses identified in this study, as well as those available in the NCBI database were calculated using SDTv1.2 (Muhire et al., 2014) based on MAFFT 7.505 (https://mafft.cbrc.jp/alignment/software) alignments with standard parameters. Virus names, abbreviations and NCBI accession numbers of ophioviruses already reported are shown in **Supplementary Table 1**.

#### 2.3.3 Phylogenetic analysis

Phylogenetic analysis based on the predicted CP protein or the polymerase protein of all available ophioviruses was done using MAFFT 7.505 with multiple aa sequence alignments using G-INS-i and E-INS-i as the best-fit model, respectively. The aligned aa sequences were used as input to generate phylogenetic trees by the maximum-likelihood method with the FastTree 2.1.11 tool available at http://www.microbesonline.org/fasttree/. Local support values were calculated with the Shimodaira-Hasegawa test (SH) and 1,000 trees resamples. The capsid proteins of two selected cytorhabdoviruses (alfalfa dwarf virus YP_009177015 and lettuce necrotic yellows virus YP_425087) were used as outgroup in the CP tree. The polymerase proteins of three related and unclassified aspivirus-like viruses (nees’ pellia aspi-like virus CAH2618860, Plasmopara viticola lesion ass. mycoophiovirus 1 QJX19787, grapevine-associated serpento-like virus 1 QXN75438) were used as outgroup in the polymerase trees. To explore the potential phylogenetic co-divergence of ophioviruses with their associated host plants, plant host cladograms were generated in phyloT v.2 (https://phylot.biobyte.de/), based on NCBI hierarchical Taxonomy. Host associations were based on connections manually inferred between viral and plant phylogram and cladograms.

## 3. Results and Discussion

Known ophioviruses are agronomically relevant, including viruses generating detrimental infections and disease in crops and ornamental plants. This *statu quo* is grounded on a tradition of biased sampling oriented to virus discovery in symptomatic and economically important plants. In this scenario, ophiovirus presence is not expected in sequencing libraries of non-symptomatic vegetables; thus, they are ideal candidates to be identified through mining of publicly available metatranscriptomic data. However, in the context of massive efforts directed to virus discovery in plants, as of today, only the partial genome of just only one novel tentative ophiovirus was discovered when publicly available transcriptome datasets were mined (Sidharthan et al., 2021). Therefore, to assess whether this apparent limited ophiovirus diversity was biological or technical, we directed our efforts to specifically address ophiovirus discovery. We extensively searched for these viruses in already available plant transcriptome datasets to expand the repertoire of plant infecting ophiovirus. This *in silico* driven search resulted in the identification of virus sequence evidence of 33 novel ophioviruses. We also detected three novel variants of members of two known ophiovirus species. This substantial number of newly discovered putative ophioviruses represents a 4.5-fold increase in the known ophioviruses, which undoubtedly show the importance of data-driven virus discovery to expand our understanding on the genomic diversity and peculiarities of virus taxa, such as the ophiovirus.

In this study, 12 tentatively full-length viral genome sequences were obtained through the identification, assembly, and curation of raw NCBI-SRA reads. Additionally, five of the putative viruses had all their RNA segments annotated, while 16 had some missing, mostly derived from technical difficulties to assemble segments which are relatively at low RNA levels during infection such as RNA 1. Importantly, 85% of the identified viruses included the detection of two or more RNA segments of the virus in the same sequencing library, which improved the level of confidence in the discovery. The detected viruses were associated with 33 different plant host species (**Table 1**). The majority of the host plants were herbaceous dicots, with 20 out of 33 identified as such. The remaining hosts were herbaceous monocots, liverworts, mosses, and ferns (**Table 1**). The genomes of 14 out of 16 viruses with all RNA segments annotated had three segments, and two monocot-associated ophioviruses had four segments (**Table 1**, **Figure 1**). Three out five of those ophioviruses whose complete genome were assembled, had four RNA segments, which is also a genomic organization of the ophiovirus Mirafiori lettuce big-vein virus (MLBVV) and lettuce ring necrosis virus (LRNV) (Garcia et al., 2017) and the recently reported carrot ophiovirus 1 (Fox et al., 2022). Thus, the most frequent genomic organization found for ophioviruses consists of three RNA segments.

**Figure 1.**
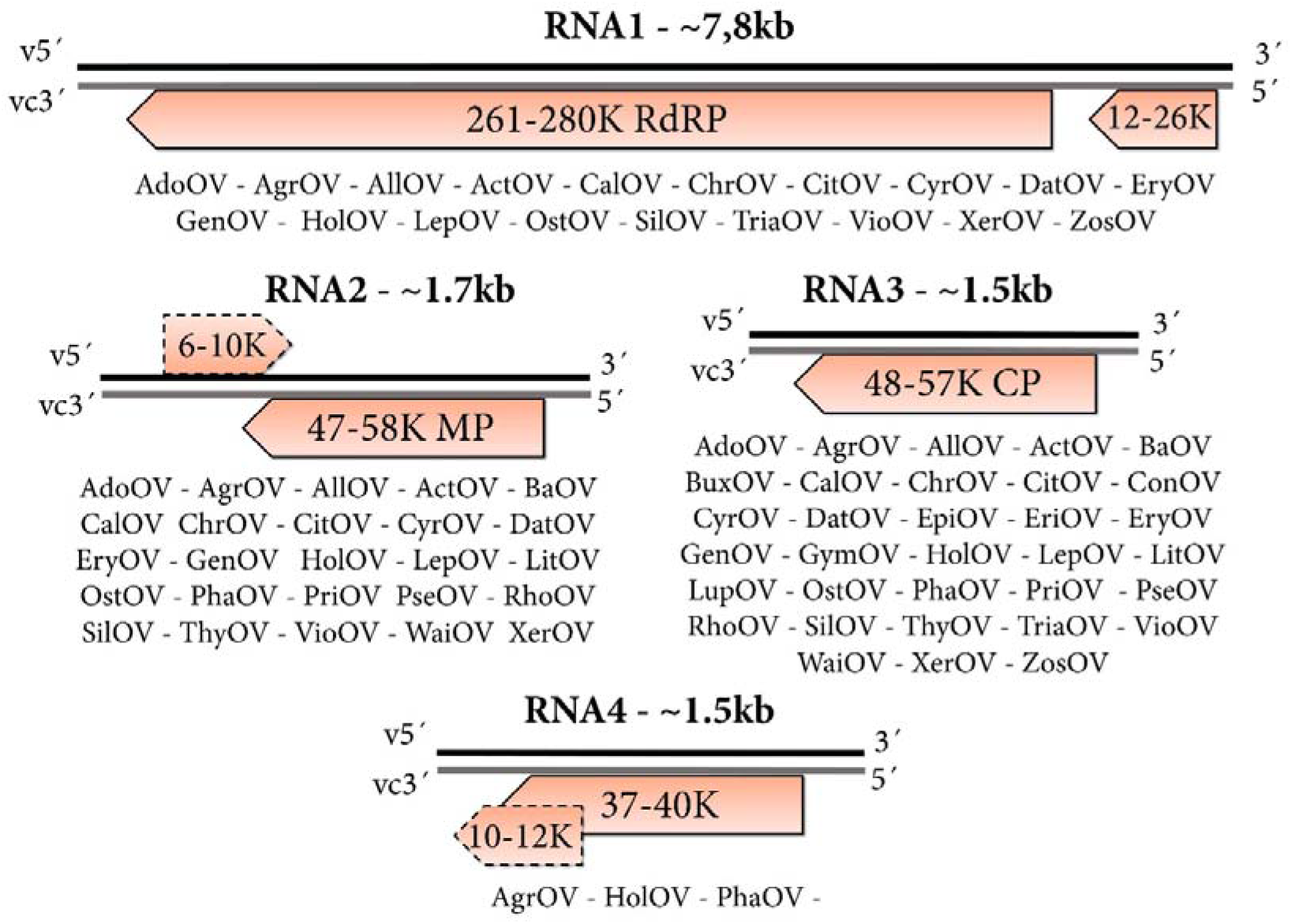
Genomic architecture of ophioviruses detected in this work. Genome graphs depicting organization and predicted gene products of each RNA segment. The predicted coding sequences are shown in orange arrowed rectangles. Dotted rectangles represent less common ORFs. Size in nucleotides and molecular weights in kilo Daltons of predicted proteins are indicated. Abbreviations: CP, capsid protein CDS; R, RNA-dependent RNA-polymerase CDS; MP, Movement Protein; v, virus RNA strand; vc, virus complementary RNA strand. Virus abbreviations are described in **Table 1**.

The RNA segments were found to encode for various proteins, including the polymerase, movement protein, and capsid protein. Like all previously reported ophioviruses (Garcia et al., 2017), the RNA 1 encoded two proteins, at 3’of the vcRNA, a large 261-280 kDa protein including the core polymerase module with the typical conserved motifs “A–E” of the RdRP, with the expected SDD signature sequence in motif “C”. Separated by an intergenic region, the other ORF at 5’of the vcRNA, encodes a small protein with a size that ranged from 105 to 245 amino acids (aa) (**Figure 1**). The RNA 1 small protein of citrus psorosis virus (CPsV), the 24K protein, has been described to localize at the nucleus, is involved in miRNA misprocessing in citrus (Reyes et al., 2016), and is an RNA silencing suppressor (Robles Luna et al., 2017). Interestingly, this small protein was quite diverse in most of the viruses identified in this study and no hits were found when a BLASTP searches were conducted (**Table 1**). The vcRNA 2 encodes a putative movement protein (MP) ranging from 47 to 58 kDa, which was characterized as a cell-to-cell MP for CPsV (54K protein) and MLBVV (55K protein) (Robles Luna et al., 2013, Hiraguri et al., 2013). All the predicted MP proteins presented the 30K core MP domain including the signature aspartate involved in cell-to-cell movement (Borniego et al., 2016). In addition, a few detected viruses encoded a small 6-10 kDa protein in the vRNA 2 with no blast hits or conserved domains, which is consistent with the proposed ambisense nature of RNA 2 postulated for MLBVV, which harbors a 10 kDa protein of unknown function at the same locus (van der Wilk et al., 2002). The vcRNA 3 encodes the capsid protein (Peña et al., 2012, Garcia et al., 2017) ranging from 48-57 kDa (**Figure 1**), and no additional ORFs. The RNA 4, which we identified only in three monocot-associated viruses, encodes a protein with unknown function with a size that ranged between 322 and 360 aa (**Table 1**). MLBVV RNA 4 contains a second overlapping ORF with no initiation codon and proposed to be expressed by a +1 translational frameshift, encoding a 10.6 kDa protein (van der Wilk et al., 2002). We failed to detect a similar additional overlapped ORF in the identified viruses, but we tentatively annotated a small ORF encoding a 12 kDa protein which was separated by an intergenic region at 3’of the vcRNA 4 of Agrostis ophiovirus which was conserved in the virus sequences of both plant hosts where these viruses were detected. Nuclear localization signals were also found in the polymerase, MP and CP encoded by the viruses identified in this study, similar to what was previously reported for ophiovirus species (Garcia et al., 2017). The pairwise aa sequence identities between the CP proteins of all reported ophioviruses, including those identified in this study, showed great diversity with an identity ranging from 14.2% to 98.9%, but importantly with a mean identity of only 32.1% (**Supplementary Figure 1**). This suggests that there is likely a substantial amount of undiscovered ophioviruses that may inhabit this virus space, despite the numerous viruses identified in this study. Using the species demarcation threshold of 85% aa identity of the CP (Garcia et al., 2017), all ophioviruses with complete CP coding regions assembled in this study with an identity below 85% were deemed as members of new ophiovirus species (**Supplementary Figure 2**), increasing the number of potential members of the genus by more than 4.5-fold. The genetic distance assessment was complemented with phylogenetic insights to provide evolutionary clues of the identified viruses.

Previous studies placed the ophiovirus in two distinct clades, one including a closer relationship between MLBVV and tulip mild mottle mosaic virus (TMMMV) and a separate clade conformed by blueberry mosaic associated virus (BlMaV) and CPsV. These two, are placed more distantly to the other ophioviruses suggesting that this might lead to the re-assignment of the existing species into two separate genera (Garcia et al., 2017). Phylogenetic analyses based on the deduced CP protein aa sequences of the detected viruses revealed a complex evolutionary history, showing distinctive groups and associations (**Figure 2**). On the one hand, the long branches linking BlMaV and CPsV in previous analyses (Garcia et al., 2017) undoubtedly constituted viral “dark matter”, as at least 17 new viruses expand the bounds of the viral sequence space amid these two viruses. This cluster now included a group of 11 viruses with affinities to BlMaV, six to CPsV, and a novel basal group of two viruses detected in *Asteraceae-plants* (**Figure 2**). The other clade of five viruses was expanded with two grasses viruses with affinities to LRNV, and the recently reported CaOV1 and PCaV linked to the MLBVV/TMMMV group and the freesia sneak virus (FreSV) and ranunculus white mottle virus (RWMV) group, respectively. More distantly, three small groups of viruses were found including four new viruses of orchids, and the third most basal group with very large branches of a virus associated to a poacea and another one to the aquatic plant *Zostera japonica*. Interestingly, a few years ago the first endogenous sequence of ophiovirus was detected in the genome of the related eelgrass *Zostera marina* (Marsile-Medun et al., 2018). In the genome of that plant a CP-like sequence was found, flanked by transposable elements, suggesting an ancient shared evolutionary history of eelgrass and ophioviruses, and the possibility that this group of plants might host contemporary ophioviruses, which is in line with the detected virus hosted by eelgrass in this work. Further, a novel divergent clade were found, mostly represented by viruses detected in basal plants such as mosses, liverworts and ferns, which represents the first association of ophioviruses with non-vascular plants and pteridophytes (**Figure 2**). Additional phylogenetic analyses based on the deduced polymerase protein aa sequences showed a similar evolutionary history of the corresponding viruses supporting the results based on CP assessment. For instance, ferns, mosses and liverworts associated ophioviruses clustered together both in CP and RdRP-based trees, suggesting that they share a unique evolutionary history among ophioviruses (**Supplementary Figure 3**). We generated a tanglegram to compare the virus phylogram and plant host cladogram to further explore potential virus-host relationships (**Figure 3**). This analysis showed that viruses of some clades clearly co-diverged with their hosts, including the orchid-associated virus clade and the clade of fern, mosses and liverworts viruses, suggesting a shared host-virus evolution in those groups (**Figure 3**). Nevertheless, the tanglegram topology also showed that for many of the ophioviruses, there is no apparent concordant evolutionary history with their potential plant hosts.

**Figure 2.**
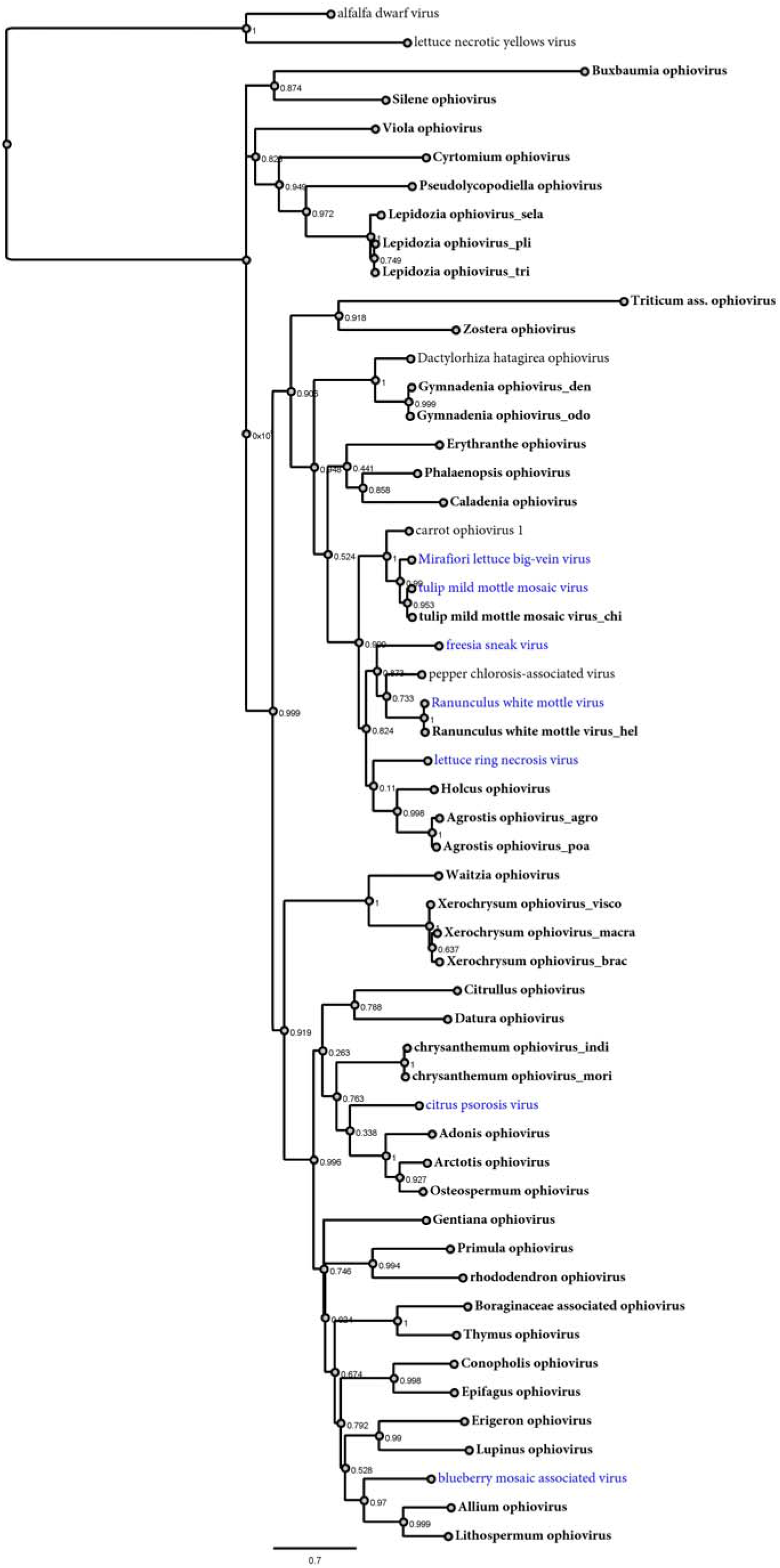
Maximum-likelihood phylogenetic tree based on the amino acid MAFFT sequence alignments of the CP protein of all the ophioviruses reported thus far and in this study. The scale bar indicates the number of substitutions per site. The node labels indicate FastTree support values. The CP proteins of two cytorhabdoviruses (alfalfa dwarf virus YP_009177015 and lettuce necrotic yellows virus YP_425087) were used as outgroups.

**Figure 3.**
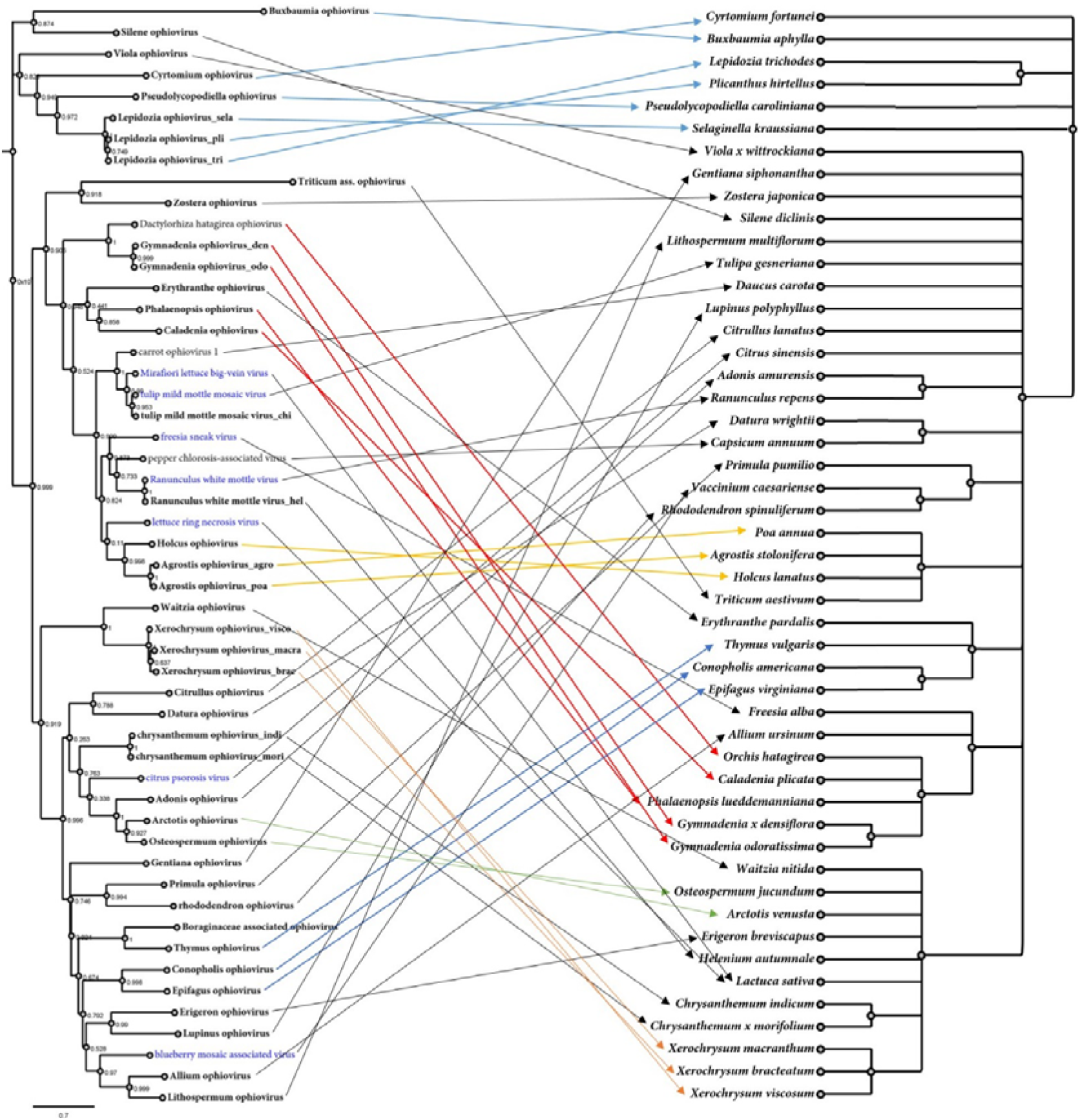
Tanglegram showing the phylogenetic relationships of the ophioviruses (left), which are linked with the associated plant host(s) shown on the right. Links of well-supported clades of viruses to taxonomically related plant species are indicated in colors. A maximum likelihood phylogenetic tree of ophioviruses was constructed based on the CP protein. Plant host cladograms were generated in phyloT v.2 based on NCBI taxonomy. Viruses identified in the present study are shown in bold font. Virus corresponding to recognized species are depicted in blue. The scale bar indicates the number of substitutions per site.

Members of four out of the seven ophiovirus species recognized so far are reported to be transmitted via soil-borne fungus of the genus *Olpidium* (Garcia et al., 2017), while CPsV, which is transmitted by vegetative propagation of the host, no natural vector had been identified (Garcia et al., 2017). Nevertheless, while we assessed in the Serratus platform thousands of sequencing libraries, we failed to robustly detect ophiovirus-like sequences in any fungal library. Interestingly, one of the ophioviruses identified in this study was discovered in a transcriptome dataset of bumblebee. Further inspection of the raw reads of this dataset retrieved a significant amount of plant reads, which based on rRNA analysis corresponded to the *Boraginacea* family. We tentatively linked this virus to this family of plants and we cautiously speculate on the possibility that this ophiovirus could be pollen-associated and transported to other plants by bumblebees. In this line, a recent study characterized the pollen virome of wild plants identifying plenty of pollen-associated viruses, but no ophiovirus (Fetter et al., 2022). Moreover, these authors found that the pollen virome is visually asymptomatic. This anecdotal observation and our difficulties in detecting ophiovirus-like sequences in fungal libraries could provide some grounds to the possibility that (some?) ophioviruses could be vertically transmitted. Other lines of evidence could support this suggestion: *i*) host-virus co-divergence in some clades may implicate isolation and a lack of horizontal transmission, *ii*) an emerging characteristic of persistent, chronic infections of several plant viruses that are vertically transmitted are latent/asymptomatic infections, a feature that could be shared by ophioviruses. Thus, further studies should be carried out to elucidate alternative transmission modes of ophioviruses beyond the fungally-transmitted MLBVV, TMMMV, LRNV and FreSV (Lot et al., 2002; Meekes and Verbeek, 2011).

There are many limitations in this study, for instance the incapacity to return to the original biological material to repeat and check the assembled viral genome sequences is a noteworthy restriction of the data mining approach for virus discovery. Similarly, contamination, spill over, and other technical artefacts could result in false positive detections or poor host assignment. In addition, a lack of a directed strategy to address virus segment termini, such as RACE, results in difficulties to determine *bona fide* RNA virus ends, which have conserved functional and structural cues in ophioviruses (Garcia et al., 2017). Some aspects of our strategy for virus discovery can overcome several of these limitations providing additional evidence on identification. For instance, the detection of the same putative virus in independent libraries from the same plant host, a robust depth coverage of virus reads, detection of more than one RNA segment of the virus in the same library, or the detection of strains of a virus in evolutionary related plants. Nevertheless, associations and detections should be complemented by further studies. All in all, based on current guidelines (Postler et al., 2022), we suggest potential latinized binomial virus species names to include the viruses described here as member of novel species within the genus *Ophiovirus* (**Table 2**). The distinctive phylogenetic clustering and the significant divergence in terms of aa identity of predicted proteins of several of the identified viruses raises questions about taxonomic classification. Currently, family *Aspiviridae* includes a single recognized genus with seven member species. Following the molecular criterion for ophiovirus species demarcation of a CP amino acid sequence identity <85%, we suggest that the identified viruses in this study could be members of novel species. In this line, it has not escaped our notice that eventually some of the groups of viruses reported here, if recognized, could be included in new genera within the *Aspiviridae* family, applying a genus demarcation criterion still not defined.

In summary, this study illustrates the significance of the analysis of NCBI-SRA public data as a valued tool not only to fasten the discovery of novel viruses but, also, to increase our understanding into their evolution and to improve virus taxonomy. Using this approach, we looked for hidden ophio-like virus sequences to expand the repertoire of these viruses expanding 4.5-fold potential existing members within the genus. Additionally, we fostered the most comprehensive phylogeny of ophioviruses to date and shed new light on the phylogenetic relationships and evolutionary landscape of this group of viruses. Future studies should focus not only on complementing our genomic predictions, but also on providing clues on the biology and ecology of these viruses such as associated symptomatology, transmission, and putative vectors.

## Acknowledgments

We would like to express genuine appreciation to the producers of the original data used for this work, which are cited in **Table 1**. By ensuing open science practices with accessible raw sequence data in open public repositories, they supported contributions based on secondary data analyses.

## Author Contributions

Conceptualization, H.D and N.B; data analysis, H.D and N.B; writing—original draft preparation, H.D and N.B; writing—review and editing, H.D, N.B, and M.L.G. All authors have read and agreed to the published version of the manuscript.

## Institutional Review Board Statement

Not applicable for studies not involving humans or animals.

## Informed Consent Statement

Not applicable for studies not involving humans.

## Data availability statement

Nucleotide sequence data reported are available in the Third Party Annotation Section of the DDBJ/ENA/GenBank databases under the accession numbers TPA: BK062646-BK062750 and can be found as **Supplementary Material 1** of this submission.

---

**Supplementary Figure 1.**
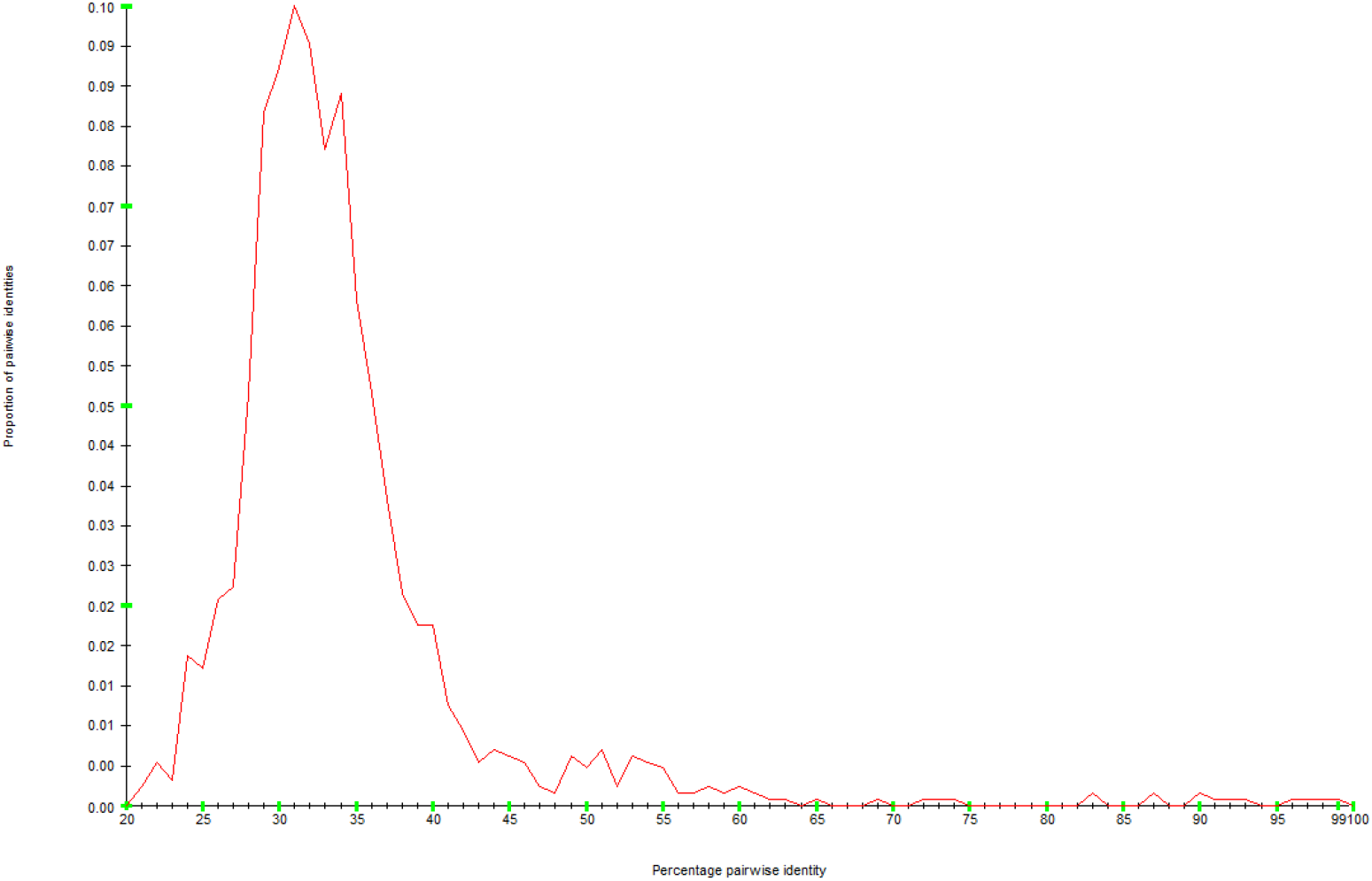
Plot of frequency of percentage pairwise identity of ophiovirus complete capsid proteins generated using SDT v1.2 software based on MAFFT amino acid sequence alignments.

**Supplementary Figure 2.**
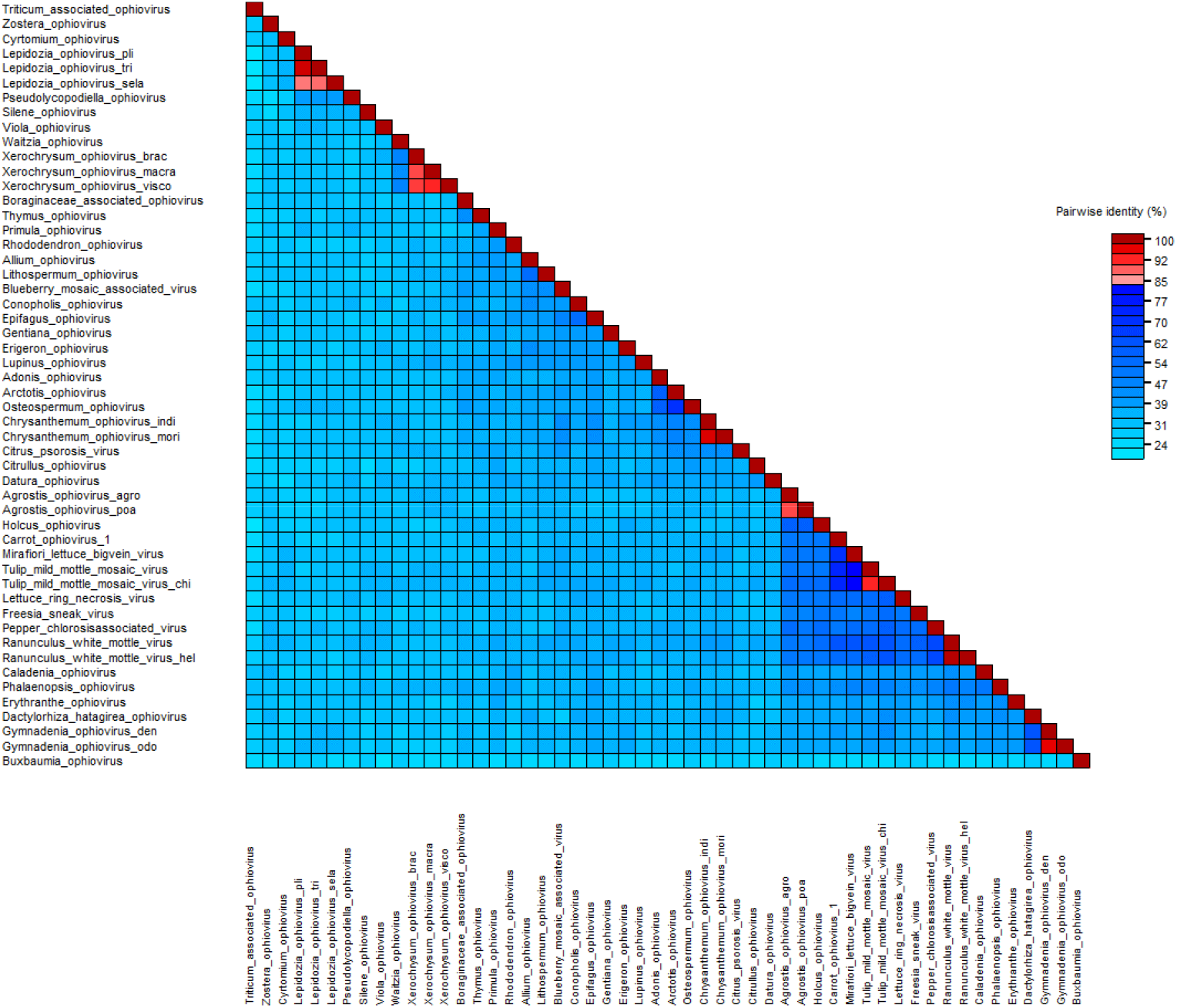
Pairwise identity matrix of the amino acid sequences of the ophiovirus complete capsid proteins generated using SDT v1.2 software based on MAFFT alignments. The colored cut-off is based on ICTV demarcation criteria of ophioviruses, which includes CP amino acid sequence identity <85% to be considered novel species (blue-light blue).

**Supplementary Figure 3.**
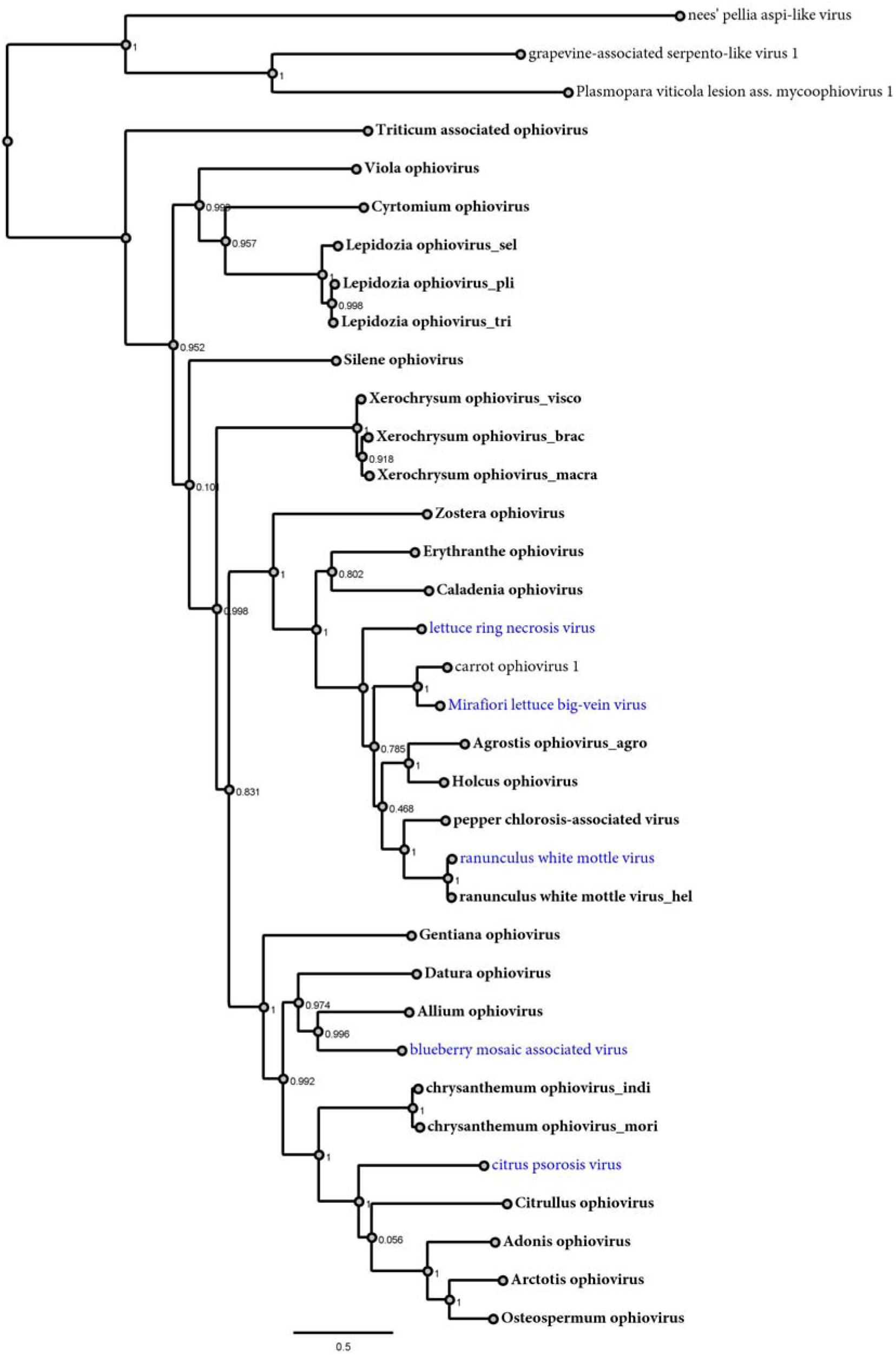
Maximum-likelihood phylogenetic tree based on the amino acid MAFFT sequence alignments of the L protein of all the ophioviruses reported thus far and in this study. The scale bar indicates the number of substitutions per site. The node labels indicate FastTree support values. The polymerase proteins of three related and unclassified aspivirus-like viruses (nees’ pellia aspi-like virus CAH2618860, Plasmopara viticola lesion ass. mycoophiovirus 1 QJX19787, grapevine-associated serpento-like virus 1 QXN75438) were used as outgroup.

## References

- Bejerman, N., & Debat, H. (2022). Exploring the tymovirales landscape through metatranscriptomics data. Archives of Virology, 167(9), 1785–1803. https://doi.org/10.1007/s00705-022-05493-9

- Bejerman, N., Debat, H., & Dietzgen, R. G. (2020). The plant negative-sense RNA virosphere: virus discovery through new eyes. Frontiers in Microbiology, 11, 588427. https://doi.org/10.3389/fmicb.2020.588427

- Bejerman, N., Dietzgen, R. G., & Debat, H. (2022). Unlocking the hidden genetic diversity of varicosaviruses, the neglected plant rhabdoviruses. Pathogens, 11(10), 1127. https://doi.org/10.3390/pathogens11101127

- Borniego, M. B., Karlin, D., Peña, E. J., Luna, G. R., & García, M. L. (2016). Bioinformatic and mutational analysis of ophiovirus movement proteins, belonging to the 30K superfamily. Virology, 498, 172–180. https://doi.org/10.1016/j.virol.2016.08.027

- Dolja, V. V., Krupovic, M., & Koonin, E. V. (2020). Deep roots and splendid boughs of the global plant virome. Annu. Rev. Phytopathol, 58(10.1146). https://doi.org/10.1146/annurev-phyto-030320-041346

- Edgar, R. C., Taylor, J., Lin, V., Altman, T., Barbera, P., Meleshko, D., … & Babaian, A. (2022). Petabase-scale sequence alignment catalyses viral discovery. Nature, 602(7895), 142–147. https://doi.org/10.1038/s41586-021-04332-2

- Fox, A., Gibbs, A. J., Fowkes, A. R., Pufal, H., McGreig, S., Jones, R. A., … & Adams, I. P. (2022). Enhanced Apiaceous Potyvirus Phylogeny, Novel Viruses, and New Country and Host Records from Sequencing Apiaceae Samples. Plants, 11(15), 1951. https://doi.org/10.3390/plants11151951

- García, M. L., Dal Bó, E., Da Graça, J. V., Gago-Zachert, S., Hammond, J., Moreno, P., … & Consortium, I. R. (2017). ICTV virus taxonomy profile: Ophioviridae. The Journal of general virology, 98(6), 1161. https://doi.org/10.1099/jgv.0.000836

- Geoghegan, J. L., & Holmes, E. C. (2017). Predicting virus emergence amid evolutionary noise. Open biology, 7(10), 170189. https://doi.org/10.1098/rsob.170189

- Hiraguri, A., Ueki, S., Kondo, H., Nomiyama, K., Shimizu, T., Ichiki-Uehara, T., … & Sasaya, T. (2013). Identification of a movement protein of Mirafiori lettuce big-vein ophiovirus. Journal of General Virology, 94(5), 1145–1150. https://doi.org/10.1099/vir.0.050005-0

- Koonin, E. V., Dolja, V. V., Krupovic, M., Varsani, A., Wolf, Y. I., Yutin, N., … & Kuhn, J. H. (2020). Global organization and proposed megataxonomy of the virus world. Microbiology and molecular biology reviews, 84(2), e00061–19. https://doi.org/10.1128/FMMBR.00061-19

- Lauber, C., & Seitz, S. (2022). Opportunities and challenges of data-driven virus discovery. Biomolecules, 12(8), 1073. https://doi.org/10.3390/biom12081073

- Leebens-Mack, J. H., Barker, M. S., Carpenter, E. J., Deyholos, M. K., Gitzendanner, M. A., Graham, S. W., … & Szövényi, P. (2019). One thousand plant transcriptomes and the phylogenomics of green plants. Nature, 574(7780), 679–685. https://doi.org/10.1038/s41586-019-1693-2

- Lot, H., Campbell, R. N., Souche, S., Milne, R. G., & Roggero, P. (2002). Transmission by Olpidium brassicae of Mirafiori lettuce virus and Lettuce big-vein virus, and their roles in lettuce big-vein etiology. Phytopathology, 92(3), 288–293. https://doi.org/10.1094/PHYTO.2002.92.3.288

- Marsile-Medun, S., Debat, H. J., & Gifford, R. J. (2018). Identification of the first endogenous Ophiovirus sequence. bioRxiv, 235044. https://doi.org/10.1101/235044

- Meekes, E. T. M., & Verbeek, M. (2011). New insights in Freesia leaf necrosis disease. Acta Horticulturae, 901, 231–236. https://doi.org/10.17660/ActaHortic.2011.901.29

- Mifsud, J. C., Gallagher, R. V., Holmes, E. C., & Geoghegan, J. L. (2022). Transcriptome Mining Expands Knowledge of RNA Viruses across the Plant Kingdom. Journal of Virology, e00260–22. https://doi.org/10.1128/jvi.00260-22

- Muhire, B. M., Varsani, A., & Martin, D. P. (2014). SDT: a virus classification tool based on pairwise sequence alignment and identity calculation. PloS one, 9(9), e108277. https://doi.org/10.1371/journal.pone.0108277

- Peña, E. J., Luna, G. R., Zanek, M. C., Borniego, M. B., Reyes, C. A., Heinlein, M., & García, M. L. (2012). Citrus psorosis and Mirafiori lettuce big-vein ophiovirus coat proteins localize to the cytoplasm and self interact in vivo. Virus research, 170(1-2), 34–43. https://doi.org/10.1016/j.virusres.2012.08.005

- Postler, T. S., Rubino, L., Adriaenssens, E. M., Dutilh, B. E., Harrach, B., Junglen, S., … & Stockdale, S. R. (2022). Guidance for creating individual and batch latinized binomial virus species names. Journal of General Virology, 103(12), 001800. https://doi.org/10.1099/jgv.0.001800

- Reyes, C. A., Ocolotobiche, E. E., Marmisollé, F. E., Robles Luna, G., Borniego, M. B., Bazzini, A. A., … & García, M. L. (2016). Citrus psorosis virus 24 K protein interacts with citrus miRNA precursors, affects their processing and subsequent miRNA accumulation and target expression. Molecular plant pathology, 17(3), 317–329. https://doi.org/10.1111/mpp.12282

- Robles Luna, G., Peña, E. J., Borniego, M. B., Heinlein, M., & Garcia, M. L. (2013). Ophioviruses CPsV and MiLBVV movement protein is encoded in RNA 2 and interacts with the coat protein. Virology, 441(2), 152–161. https://doi.org/10.1016/j.virol.2013.03.019

- Robles Luna, G., Reyes, C. A., Peña, E. J., Ocolotobiche, E., Baeza, C., Borniego, M. B., … & García, M. L. (2017). Identification and characterization of two RNA silencing suppressors encoded by ophioviruses. Virus research, 235, 96–105. https://doi.org/10.1016/j.virusres.2017.04.013

- Shimomoto, Y., Takemura, C., Yanagisawa, H., Neriya, Y., & Sasaya, T. (2023). Complete genome sequence of a novel ophiovirus associated with chlorotic disease of pepper (Capsicum annuum L.) in Japan. Archives of Virology, 168(2), 48. https://doi.org/10.1007/s00705-022-05691-5

- Sidharthan, V. K., Kalaivanan, N. S., & Baranwal, V. K. (2021). Discovery of putative novel viruses in the transcriptomes of endangered plant species native to India and China. Gene, 786, 145626. https://doi.org/10.1016/j.gene.2021.145626

- Simmonds, P., Adams, M. J., Benkő, M., Breitbart, M., Brister, J. R., Carstens, E. B., … & Zerbini, F. M. (2017). Virus taxonomy in the age of metagenomics. Nature Reviews Microbiology, 15(3), 161–168. https://doi.org/10.1038/nrmicro.2016.177

- Van der Wilk, F., Dullemans, A. M., Verbeek, M., & Van Den Heuvel, J. F. J. M. (2002). Nucleotide sequence and genomic organization of an ophiovirus associated with lettuce big-vein disease. Journal of General Virology, 83(11), 2869–2877. https://doi.org/10.1099/0022-1317-83-11-2869

