## Supplementary Table 1 for "Expanding the repertoire of the plant infecting ophioviruses"

Supplementary Table S1. Virus names, abbreviations and NCBI accession numbers of ophiovirus sequences used in this study

| **Virus name** | **Abbreviation** | **Accession number** |
| --- | --- | --- |
| blueberry mosaic associated virus | BlMaV | KJ704366- KJ704368 |
| carrot ophiovirus 1 | CaOV1 | OM419178-OM419181 |
| citrus psorosis virus | CPsV | MZ330078-MZ330080 |
| Dactylorhiza hatagirea ophiovirus | DhOV | BK013328 |
| freesia sneak virus | FreSV | MN365016 |
| lettuce ring necrosis virus | LRNV | MW594439- MW594442 |
| Mirafiori lettuce big-vein virus | MLBVV | AF525933- AF525936 |
| ranunculus white mottle virus | RWMV | KY769680 |
| tulip mild mottle mosaic virus | TMMMV | KY769675 |
| pepper chlorosis-associated virus | PCaV | BDO47352-BDO47349 |
