## Supplementary material for "Expanding the repertoire of the plant infecting ophioviruses": Table 1

**Table 1**. Summary of novel ophioviruses identified from plant RNA-seq data available in the NCBI database. Acronyms of best hits are listed in Supplementary Table 1.

| **Plant host** | **Taxa/**  **family** | **Virus name/**  **Abbreviation** | **Bioproject ID/**  **Data citation** | **Segment** | **Length (nt)** | **Accession number** | **Protein ID** | **Length (aa)** | **Highest scoring virus- protein/*E*-value/query coverage%/identity% (Blast P)** |
| --- | --- | --- | --- | --- | --- | --- | --- | --- | --- |
| Amur adonis  (*Adonis amurensis*) | dicot/  *Ranunculaceae* | Adonis ophiovirus/  AdoOV | PRJNA521968/  Zhou et al., (2019) | RNA1  RNA2  RNA3 | 7425*  1595  1448 | BK062646  BK062647  BK062648 | L  MP  CP | 2411*  467  450 | CPsV-L/0.0/93/46.79  CPsV-MP/3e-108/94/42.07  CPsV-CP/6e-110/100/40.44 |
| Creeping bentgrass  (*Agrostis stolonifera*) | monocot/  *Poaceae* | Agrostis ophiovirus_agro/  AgrOV_agro | PRJNA324407/  Ma et al., (2017) | RNA1  RNA2  RNA3  RNA4 | 7710*  1863  1540  1907 | BK062649  BK062650  BK062651  BK062652 | L  22kDa  MP  CP  37kDa | 2294*  174  499  453  322 | MLBVV-L/0.0/84/59.61  RWMV-22kDa/1e-21/75/40.91  MLBVV-MP/1e-155/98/49  MLBVV-CP/3e-138/99/49  LRNV-37kDa/4e-69/90/37.54 |
| Annual bluegrass  (*Poa annua*) | monocot/  *Poaceae* | Agrostis ophiovirus_poa/  AgrOV_poa | PRJNA265116/  Chen et al., (2016) | RNA 1  RNA2  RNA3  RNA4 | 824*  1828  1525  1428 | BK062653  BK062654  BK062655  BK062656 | 22kDa  MP  CP  37kDa | 174  499  453  323 | RWMV-22kDa/7e-21/77/40.96  MLBVV-MP/5e-155/98/49  MLBVV-CP/1e-138/99/49  LRNV-37kDa/7e-69/90/37.59 |
| Wild garlic  (*Allium ursinum*) | monocot/  *Amaryllidaceae* | Allium ophiovirus/  AllOV | PRJNA542932/  Fakjus et al., (2019) | RNA1  RNA2  RNA3 | 7380*  1832  1495 | BK062657  BK062658  BK062659 | L  MP  CP | 2338  478  454 | BlMaV-L/0.0/99/52.96  BlMaV-MP/4e-111/93/38  BlMaV-CP/5e-125/81/49.6 |
| Silver actotis  (*Arctotis venusta*) | dicot/  *Asteraceae* | Arctotis ophiovirus/  ActOP | PRJNA371565/  Jayasena et al., (2017) | RNA1  RNA2  RNA3 | 8319  1738  1462 | BK062660  BK062661  BK062662 | L  22kDa  MP  CP | 2406  222  486  446 | CPsV-L/0.0/99/45.18  no hits  CPsV-MP/2e-142/97/48  CPsV-CP/1e-121/100/43.56 |
| Borage  (*Boranginaceae*) | dicot/  *Boranginaceae* | Boranginaceae associated ophiovirus/BaOV | PRJNA659133/  Sun et al., 2021 | RNA2  RNA3 | 1737  1589 | BK062663  BK062664 | MP  CP | 486  471 | BlMaV-MP/2e-110/91/39.74  BlMaV-CP/2e-86/76/41.92 |
| Bug moss  (*Buxbaumia aphylla*) | Bryophyta/  *Buxbaumiaceae* | Buxbaumia ophiovirus/  BuxOV | PRJEB21674/  1000 Plant (1KP) Transcriptomes Initiative | RNA3 | 1590 | BK062665 | CP | 485 | MLBVV-CP/4e-18/53/25.82 |
| Crab-lipped spider orchid (*Caladenia plicata*) | monocot/  *Orchidaceae* | Caladenia ophiovirus/  CalOV | PRJNA384875/  [Xu et al., 2017] | RNA1  RNA2  RNA3 | 7488  1760  1423 | BK062666  BK062667  BK062668 | L  22kDa  MP  CP | 2247  163  445  438 | RWMV-L/0.0/100/49.48  RWMV-22kDa/7e-10/61/32.67  LRNV-MP/9e-100/99/39.47  RWMV-CP/4e-101/94/39.66 |
| Indian chrysanthemum  (*Chrysanthemum indicum*) | dicot/  *Asteraceae* | Chrysanthemum ophiovirus_indi/ChrOV_indi | PRJNA361213/  Han et al., (2018) | RNA1  RNA2  RNA3 | 8240  2143  1572 | BK062669  BK062670  BK062671 | L  22kDa  MP  CP | 2379  222  483  457 | BlMaV-L/0.0/97/47.45  BlMaV-22kDa/0.002/62/30.54  CPsV-MP/4e-117/94/42.61  BlMaV-CP/3e-109/99/42.19 |
| Garden mum  (*Chrysanthemum morifolium*) | dicot/  *Asteraceae* | Chrysanthemum ophiovirus_mori/ChrOV_mori | PRJNA315793/  Zhang et al., (2015) | RNA1  RNA2  RNA3 | 8255  2164  1573 | BK062672  BK062663  BK062674 | L  22kDa  MP  CP | 2379  222  483  457 | BlMaV-L/0.0/97/47.49  BlMaV-22kDa/0.002/62/30.50  CPsV-MP/4e-117/94/42.64  BlMaV-CP/3e-109/99/42.15 |
| Watermelon  (*Citrullus lanatus*) | dicot/  *Cucurbitaceae* | Citrullus ophiovirus/CitOV | PRJNA576654/  Garcia-Lozano et al., (2019) | RNA1  RNA2  RNA3 | 8510  1760  1528 | BK062675  BK062676  BK062677 | L  22kDa  MP  CP | 2418  245  483  464 | CPsV-L/0.0/99/42.77  no hits  CPsV-MP/9e-127/97/43.19  CPsV-CP/8e-82/93/38.66 |
| Bear corn  (*Conopholis americana*) | dicot/  *Orobanchaceae* | Conopholis ophiovirus/ConOV | PRJEB21674/  1000 Plant (1KP) Transcriptomes Initiative | RNA3 | 1684 | BK062678 | CP | 481 | BlMaV-CP/4e-99/69/45.35 |
| Holly fern  (*Cyrtomium fortunei*) | *Polypodiophyta/*  *Dryopteridaceae* | Cyrtomium ophiovirus/CyrOV | PRJNA384992/  You et al., (2017) | RNA1  RNA2  RNA3 | 7548  1902  1749 | BK062679  BK062680  BK062681 | L  22kDa  MP  CP | 2357  105  409  500 | BlMaV-L/0.0/97/36.83  no hits  BlMaV-MP/4e-06/67/22.92  MLBVV-CP/4e-50/67/33.90 |
| Sacred datura  (*Datura wrightii*) | Dicot/  *Solanaceae* | Datura ophiovirus/DatOV | PRJNA473174/  Sun, University of California, USA | RNA1  RNA2  RNA3 | 8055  1788  1701 | BK062682  BK062683  BK062684 | L  22kDa  MP  CP | 2366  186  481  511 | BlMaV-L/0.0/97/50.09  no hits  BlMaV-MP/1e-86/95/35.43  CPsV-CP/1e-88/69/39.50 |
| Beech drops  (*Epifagus virginiana*) | dicot/  *Orobanchaceae* | Epifagus ophiovirus/EpiOV | PRJEB21674/  1000 Plant (1KP) Transcriptomes Initiative | RNA3 | 1371* | BK062685 | CP | 328* | BlMaV-CP/3e-59/88/42.81 |
| Lifeflower  (*Erigeron breviscapus*) | dicot/  *Asteraceae* | Erigeron ophiovirus/EriOV | PRJNA293262/  Chen et al., (2015) | RNA3 | 1837 | BK062686 | CP | 463 | BlMaV-CP/2e-92/74/44.09 |
| Pardus monkey-flower  (*Erythranthe* *pardalis*) | dicot/  *Phrymaceae* | Erythranthe ophiovirus/EryOV | PRJNA508749/  Flores-Vergara et al., (2020) | RNA1  RNA2  RNA3 | 7643  1587  1651 | BK062687  BK062688  BK062689 | L  22kDa  MP  CP | 2271  195  436  490 | LRNV-L/0.0/99/51.73  BlMaV-22kDa/2e-11/58/30.43  LRNV-MP/1e-97/90/39.33  RWMV-CP/1e-100/99/36.68 |
| Tube gentian  (*Gentiana siphonantha*) | dicot/  *Gentianaceae* | Gentiana ophiovirus/ (GenOV) | PRJNA555883/  Chen et al., (2021) | RNA1  RNA2  RNA3 | 8043  2077  1473 | BK062690  BK062691  BK062692 | L  22kDa  MP  CP | 2254  190  516  450 | BlMaV-L/0.0/99/45.32  BlMaV-22kDa/0.007/75/24.32  BlMaV-MP/3e-47/52/37.86  BlMaV-CP/2e-92/80/42.66 |
| Marsh fragrant orchid  (*Gymnadenia densiflora*) | monocot/  *Orchidaceae* | Gymnadenia ophiovirus_den/GymOV_den | PRJNA504609/  Piñeiro Fernandez et al., (2019) | RNA3 | 1431 | BK062693 | CP | 446 | DhOV-CP/0.0/94/60.05 |
| Short-spurred fragrant orchid  (*Gymnadenia odorattissima*) | monocot/  *Orchidaceae* | Gymnadenia ophiovirus_odo/GymOV_odo | PRJNA504609/  Piñeiro Fernandez et al., (2019) | RNA3 | 1339* | BK062694 | CP | 425 | DhOV-CP/0.0/87/58.25 |
| Common velvetgrass  (*Holcus lanatus*) | monocot/  *Poaceae* | Holcus ophiovirus/HolOV | PRJEB3994/  Young et al., (2018) | RNA 1  RNA2  RNA3  RNA4 | 7627*  1770  1495  1436 | BK062695  BK062696  BK062697  BK062698 | L  22kDa  MP  CP  37kDa | 2194*  162  459  444  322 | RWMV-L/0.0/89/65.13  MLBVV-22kDa/3e-24/82/40  MLBVV-MP/2e-176/99/53.43  RWMV-CP/4e-148/100/49.77  LRNV-37kDa/1e-73/98/39.06 |
| Hairy liverwort  (*Lepidozia trichodes*) | *Marchantiophyta*/  *Lepidoziaceae* | Lepidozia ophiovirus_tri/LepOV_tri | PRJNA505755/  Fairylake Botanical Garden, China | RNA1  RNA2  RNA3 | 7644  1872  1581 | BK062699  BK062700  BK062701 | L  22kDa  MP  CP | 2357  109  460  471 | BlMaV-L/0.0/96/37.15  no hits  BlMaV-MP/3e-19/49/25.96  MLBVV-CP/6e-58/71/30.99 |
| Basket liverwort  (*Plicanthus hirtellus*) | *Marchantiophyta*/  *Anastrophyllaceae* | Lepidozia ophiovirus_pli/LepOV_pli | PRJNA505755/  Fairylake Botanical Garden, China | RNA1  RNA2  RNA3 | 7546  1497  1555 | BK062702  BK062703  BK062704 | L  22kDa  MP  CP | 2357  109  460  471 | BlMaV-L/0.0/96/37.19  no hits  BlMaV-MP/2e-19/49/25.94  MLBVV-CP/8e-58/71/30.92 |
| Krauss' spike moss  (*Selaginella kraussiana*) | Lycophyta/  *Selaginellaceae* | Lepidozia ophiovirus_sela/LepOV_sela | PRJNA351923/  James et al., (2017) | RNA1  RNA2  RNA3 | 7644  1872  1581 | BK062705  BK062706  BK062707 | L  22kDa  MP  CP | 2357  109  460  471 | BlMaV-L/0.0/96/37.11  no hits  BlMaV-MP/4e-19/49/25.99  MLBVV-CP/5e-58/71/30.95 |
| Manyflowered gromwell  (*Lithospermum multiflorum*) | dicot/  *Boraginaceae* | Lithospermum ophiovirus/LitOV | PRJNA353131/  Cohen et al., (2016) | RNA2  RNA3 | 1498*  1693 | BK062708  BK062709 | MP  CP | 470*  460 | BlMaV-MP/7e-118/92/42.61  MLBVV-CP/1e-51/64/32.70 |
| Garden lupin  (*Lupinus polyphyllus*) | dicot/  *Fabaceae* | Lupinus ophiovirus/LupOV | PRJEB8056/  Cannon et al., (2015) | RNA3 | 1838 | BK062710 | CP | 448 | BlMaV-CP/1e-95/73/44.31 |
| Trailing Pink Daisy (*Osteospermum jucundum*) | dicot/  *Asteraceae* | Osteospermum ophiovirus/OstOV | PRJNA371565/  Jayasena et al., (2017) | RNA1  RNA2  RNA3 | 8521  1701  1512 | BK062711  BK062712  BK062713 | L  22kDa  MP  CP | 2407  204  482  449 | CPsV-L/0.0/100/46.10  no hits  CPsV-MP/2e-143/100/47.89  CPsV-CP/2e-120/100/43.65 |
| Moth orchid  (*Phalaenopsis lueddemanniana*) | monocot/  *Orchidaceae* | Phalaenopsis ophiovirus/PhaOV | PRJNA345261/  Chao et al., (2017) | RNA2  RNA3  RNA4 | 1867  1630  1296 | BK062714  BK062715  BK062716 | MP  CP  37kDa | 489  431  360 | LRNV-MP/1e-92/95/37.58  MLBVV-CP/7e-120/100/44.87  LRNV-37kDa/1e-14/47/31.76 |
| Clammy primrose  (*Primula pumilio*) | dicot/  *Primulaceae* | Primula ophiovirus/PriOV | PRJNA544345/  Fu et al., (2019) | RNA2  RNA3 | 1565  1572 | BK062717  BK062718 | MP  CP | 450  455 | LRNV-MP/2e-56/91/30.64  CPsV-CP/1e-86/93/37.15 |
| Slender bog club-moss  (*Pseudolycopodiella caroliniana*) | Lycophyta/  *Lycopodiaceae* | Pseudolycopodiella ophiovirus/PseOV | PRJEB4921/  1000 Plant (1KP) Transcriptomes Initiative | RNA2  RNA3 | 1829  1594 | BK062719  BK062720 | MP  CP | 464  466 | MLBVV-MP/1e-22/57/28.37  MLBVV-CP/2e-55/71/31.86 |
| Firecracker rhododendron  (*Rhododendron spinuliferum*) | dicot/  *Ericaceae* | Rhododendron ophiovirus/RhoOV | PRJNA530078/  Xue Zhang, Yunnan University, China | RNA2  RNA3 | 1867  1406 | BK062725  BK062726 | MP  CP | 452  441 | BlMaV-MP/3e-49/95/31.25  BlMaV-CP/6e-78/95/37.07 |
| Diclinis campion  (*Silene diclinis*) | dicot/  *Caryophyllaceae* | Silene ophiovirus/  SilOV | PRJEB39526/  Muyle et al., (2021) | RNA1  RNA2  RNA3 | 6036*  1511  1532 | BK062727  BK062728  BK062729 | L  MP  CP | 1993*  426  446 | BlMaV-L/0.0/98/40.51  LRNV-MP/2e-29/69/29.30  TMMMV-CP/2e-53/80/35.12 |
| Thyme  (*Thymus vulgaris*) | dicot/  *Lamiaceae* | Thymus ophiovirus/ ThyOV | PRJNA417241/  Mollion et al., (2018) | RNA2  RNA3 | 1598  1589 | BK062730  BK062731 | MP  CP | 480  477 | BlMaV-MP/3e-114/95/40.47  BlMaV-CP/1e-91/71/42.98 |
| Wheat  (*Triticum aestivum*) | monocot/  *Poaceae* | Triticum associated ophiovirus/TriaOV | PRJNA432496/  Iquebal et al., (2019) | RNA1  RNA3 | 5377*  1192* | BK062733  BK062734 | L  CP | 1792*  377* | RWMV-L/0.0/95/34.62  DhOV-CP/8e-15/57/30.77 |
| Pansies  (*Viola x wittrockiana*) | dicot/  Violaceae | Viola ophiovirus/VioOV | PRJNA552204/  Du et al., (2019) | RNA1  RNA2  RNA3 | 7671  1570  1576 | BK062735  BK062736  BK062737 | L  22kDa  MP  CP | 2308  173  435  492 | MLBVV-L/0.0/94/37.76  no hits  BlMaV-MP/4e-16/54/27.17  MLBVV-CP/4e-52/69/33.62 |
| Golden waitzia  (*Waitzia nitida*) | dicot/  *Asteraceae* | Waitzia ophiovirus /(WaiOV) | PRJNA371565/  Jayasena et al., (2017) | RNA2  RNA3 | 1570  1486 | BK062738  BK062739 | MP  CP | 453  460 | BlMaV-MP/5e-35/83/29.84  CPsV-CP/7e-63/97/31.32 |
| Strawflower  (*Xerochrysum bracteatum*) | dicot/  *Asteraceae* | Xerochrysum ophiovirus_brac/ XerOV_brac_ | PRJNA371565/  Jayasena et al., (2017) | RNA1  RNA2  RNA3 | 7681  1577  1570 | BK062740  BK062741  BK062742 | L  22kDa  MP  CP | 2266  199  444  461 | BlMaV-L/0.0/99/40.75  no hits  LRNV-MP/4e-39/90/28.76  CPsV-CP/7e-56/94/30.16 |
| White strawflower  (*Xerochrysum macranthum*) | dicot/  *Asteraceae* | Xerochrysum ophiovirus_macra/ XerOV_macra | PRJNA371565/  Jayasena et al., (2017) | RNA1  RNA2  RNA3 | 7692  1646  1513 | BK062743  BK062744  BK062745 | L  22kDa  MP  CP | 2264  199  444  459 | BlMaV-L/0.0/99/40.75  no hits  LRNV-MP/4e-40/90/29.82  CPsV-CP/7e-57/95/30.11 |
| Sticky everlasting  (*Xerochrysum viscosum*) | dicot/  *Asteraceae* | Xerochrysum ophiovirus_visco/ XerOV_visco | PRJNA371565/  Jayasena et al., (2017) | RNA1  RNA2  RNA3 | 7591  1577  1522 | BK062746  BK062747  BK062748 | L  22kDa  MP  CP | 2266  199  441  459 | BlMaV-L/0.0/99/41.45  no hits  LRNV-MP/2e-41/91/30.05  CPsV-CP/3e-58/96/30.44 |
| Dwarf eelgrass  (*Zostera japonica*) | monocot/  *Zosteraceae* | Zostera ophiovirus/  ZosOV | PRJNA419030/  Crump et al., (2018) | RNA1  RNA3 | 7748  1634 | BK062749  BK062750 | L  22kDa  CP | 2281  216  452 | LRNV-L/0.0/93/41.84  no hits  RWMV-CP/4e-52/99/30.31 |

*partial sequence
