## Supplementary material for "Expanding the repertoire of the plant infecting ophioviruses": Table 2

**Table 2**. Novel viruses: virus name and tentative species names within genus *Ophiovirus*.

| **Virus name/Abbreviation** | **Species name** |
| --- | --- |
| Adonis ophiovirus/AdoOV | *Ophiovirus adonidis* |
| Agrostis ophiovirus_agro/AgrOV_agro | *Ophiovirus agrostis* |
| Agrostis ophiovirus_poa/AgrOV_poa | *Ophiovirus agrostis* |
| Allium ophiovirus/AllOV | *Ophiovirus alli* |
| Arctotis ophiovirus/ActOP | *Ophiovirus arctotis* |
| Boranginaceae associated ophiovirus/BaOV | *Ophiovirus boranginaceae* |
| Buxbaumia ophiovirus/BuxOV | *Ophiovirus buxbaumiae* |
| Caladenia ophiovirus/CalOV | *Ophiovirus caladeniae* |
| chrysanthemum ophiovirus_indi/ChrOV_indi | *Ophiovirus chrysanthemi* |
| chrysanthemum ophiovirus_mori/ChrOV_mori | *Ophiovirus* *chrysanthemi* |
| Citrullus ophiovirus/CitOV | *Ophiovirus citrullus* |
| Conopholis ophiovirus/ConOV | *Ophiovirus conopholis* |
| Cyrtomium ophiovirus/CyrOV | *Ophiovirus cyrtomii* |
| Datura ophiovirus/DatOV | *Ophiovirus daturi* |
| Epifagus ophiovirus/EpiOV | *Ophiovirus epifagus* |
| Erigeron ophiovirus/EriOV | *Ophiovirus erigeron* |
| Erythranthe ophiovirus/EryOV | *Ophiovirus erythranthis* |
| Gentiana ophiovirus/ (GenOV) | *Ophiovirus gentianae* |
| Gymnadenia ophiovirus_den/GymOV_den | *Ophiovirus gymnadeniae* |
| Gymnadenia ophiovirus_odo/GymOV_odo | *Ophiovirus gymnadeniae* |
| Holcus ophiovirus/HolOV | *Ophiovirus holci* |
| Lepidozia ophiovirus_tri/LepOV_tri | *Ophiovirus lepidoziae* |
| Lepidozia ophiovirus_pli/LepOV_pli | *Ophiovirus lepidoziae* |
| Lepidozia ophiovirus_sela/LepOV_sela | *Ophiovirus lepidoziae* |
| Lithospermum ophiovirus/LitOV | *Ophiovirus lithospermi* |
| Lupinus ophiovirus/LupOV | *Ophiovirus lupini* |
| Osteospermum ophiovirus/OstOV | *Ophiovirus osteospermi* |
| Phalaenopsis ophiovirus/PhaOV | *Ophiovirus phalaenopsis* |
| Primula ophiovirus/PriOV | *Ophiovirus primuli* |
| Pseudolycopodiella ophiovirus/PseOV | *Ophiovirus pseudolycopodiellae* |
| rhododendron ophiovirus/RhoOV | *Ophiovirus rhododendri* |
| Silene ophiovirus/SilOV | *Ophiovirus sileni* |
| Thymus ophiovirus/ ThyOV | *Ophiovirus thymi* |
| Triticum associated ophiovirus/TriaOV | *Ophiovirus tritici* |
| Viola ophiovirus/VioOV | *Ophiovirus violae* |
| Waitzia ophiovirus /(WaiOV) | *Ophiovirus waitziae* |
| Xerochrysum ophiovirus_brac/ XerOV_brac_ | *Ophiovirus xerochrysi* |
| Xerochrysum ophiovirus_macra/ XerOV_macra | *Ophiovirus xerochrysi* |
| Xerochrysum ophiovirus_visco/ XerOV_visco | *Ophiovirus xerochrysi* |
| Zostera ophiovirus/ZosOV | *Ophiovirus zosterae* |
